# Calculating and interpreting *F*_*ST*_ in the genomics era

**DOI:** 10.1101/2024.09.24.614506

**Authors:** Menno J. de Jong, Cock van Oosterhout, A. Rus Hoelzel, Axel Janke

## Abstract

The relative genetic distance between populations is commonly measured using the fixation index (*F*_*ST*_). Traditionally inferred from allele frequency differences, the question arises how *F*_*ST*_ can be estimated and interpreted when analysing genomic datasets with low sample sizes. Here, we advocate an elegant solution first put forward by Hudson et al. (1992): *F*_*ST*_ = (*D*_*xy*_ – *π*_*xy*_)/*D*_*xy*_, where *D*_*xy*_ and *π*_*xy*_ denote mean sequence dissimilarity *between* and *within* populations, respectively. This multi-locus *F*_*ST*_-metric can be derived from allele frequency data, but also from sequence alignment data alone, even when sample sizes are low and/or unequal. As with other *F*_*ST*_-metrices, the numerator denotes net divergence (*D*_*a*_), which is equivalent to the *f*^*2*^-statistic and Nei’s *D* (for realistic estimates of *D*_*xy*_ and *π*_*xy*_). In terms of demographic inference, net divergence measures the difference in increase of *D*_*xy*_ and *π*_*xy*_ since the population split, owing to a reduction of coalescence times within populations as a result of genetic drift. Because different combinations of *ΔD*_*xy*_ and *Δπ*_*xy*_ can produce identical *F*_*ST*_-estimates, no universal relationship exists between *F*_*ST*_ and population split time. Still, in case of recent population splits, when novel mutations are negligible, *F*_*ST*_-estimates can be accurately converted into coalescent units (*τ*. i.e., split time in multiples of 2*N*_*e*_). This then allows to quantify gene tree discordance, without the need for multispecies coalescent based analyses, using the formula: *P*_*discordance*_ = ⅔·(1 – *F*_*ST*_). To facilitate the use of the Hudson *F*_*ST*_-metric, we implemented new utilities in the R package SambaR.

## INTRODUCTION

As originally defined by Wright (1943), *F*_*ST*_ and *F*_*IS*_ indicate the reduction in heterozygosity due to population structure (i.e., Wahlund effect) and close-kin matings (i.e., inbreeding), respectively (Wright 1949). Since then, various derivations of *F*_*ST*_ have been suggested as measures of levels of population differentiation (Cockerham 1973; Nei 1973; Nei 1977; Crow and Aoki 1984; Weir and Cockerham 1984; Cockerham and Weir 1993; Meirmans and Hedrick 2011; Weir 2012; Bhatia et al. 2013; Jakobsson et al. 2013; Chen et al. 2015; Berner 2019; Ochoa and Storey 2021). A common feature is that they quantify allele frequency differences, which raises the question of how *F*_*ST*_-estimates should be inferred and understood when analysing genomes from a small number of individuals per population (Willing et al. 2012).

Here, we promote an elegant solution suggested by Hudson et al. (1992), namely: *F*_*ST*_ = (*D*_*xy*_ – *π*_*xy*_)/*D*_*xy*_. Conveniently, this Hudson *F*_*ST*_-metric can be derived from allele frequency data as well as from sequence alignment data, even when sample sizes are low and/or unequal (Bhatia et al. 2013). It can be shown that the metric is consistent with the classic *F*_*ST*_-metric of Wright (1943), that the nominator is equivalent to the *f*^*2*^-statistic (Reich et al. 2009) and Nei’s D (Nei 1972), and that the metric can be inferred from multi-locus phylogenetic trees as the proportional length of inner branches.

By rewriting *F*_*ST*_ in terms of sequence dissimilarity rather than allele frequency differences, Hudson et al. (1992) created a renewed and intuitive understanding of *F*_*ST*_, without its original meaning being lost. As the formula implies, *F*_*ST*_ measures how much smaller mean sequence dissimilarity *within* two populations (*π*_*xy*_) is compared to that *between* the two populations (*D*_*xy*_) (Takahata and Nei 1985). Given that in panmictic populations *π*_*xy*_ is expected to equal genome-wide heterozygosity (*He*, proportion of heterozygous sites in diploid genomes), Hudson *F*_*ST*_ measures the reduction of heterozygosity owing to population structure, true to its original definition.

Because *D*_*xy*_ and *π*_*xy*_ are proportional to coalescence time, the definition of Hudson et al. (1992) allows for a better understanding of *F*_*ST*_ from the perspective of coalescent theory. As we will argue, *F*_*ST*_ measures the difference in increase of *D*_*xy*_ and *π*_*xy*_ since the population split (i.e., Δ*D*_*xy*_ – *Δπ*_*xy*_). This difference arises because finite populations lose haplotypes through genetic drift, which reduces the coalescence times within populations but not the coalescence times between populations. We will also argue that in case of relatively recent population split events, *F*_*ST*_-estimates can be converted into coalescent units (*τ*, i.e., split time in multiples of 2*N*_*e*_) (Slatkin 1991), therewith bypassing the more computationally demanding multispecies coalescent-based analyses.

We aim to provide an intuitive explanation of the population-genetic theory behind *F*_*ST*_, particularly those aspects which are useful for analysing genomic data. We support our conclusions with simulation outcomes, and through analyses of a population-genomic dataset of brown bears (de Jong et al. 2023). To facilitate the use of the Hudson *F*_*ST*_-metric, we have implemented new utilities to the R package SambaR (de Jong et al. 2021).

## RESULTS AND DISCUSSION

### Calculating Hudson *F*_***ST***_

Following the definition of Hudson et al. (1992), the relative genetic distance between two populations, *X* and *Y*, can be defined and understood as how much smaller mean sequence dissimilarity is *within* two populations (nucleotide diversity, *π*_*xy*_) compared to *between* the two populations (absolute genetic distance, *D*_*xy*_):

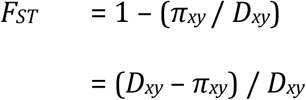

*π*_*xy*_ and *D*_*xy*_ represent here expected values (i.e., *E*(*π*_*xy*_) and *E*(*D*_*xy*_)), averaged over all sites (i.e., base pairs) in the dataset (Box 1). These sites ideally represent numerous unlinked loci, such that single-locus stochastics are cancelled out. When inferred from such a multi-locus dataset, the *F*_*ST*_-measure can be read from multi-locus trees as the proportional length of inner branches (Fig. 1A-B).

**Figure 1.**
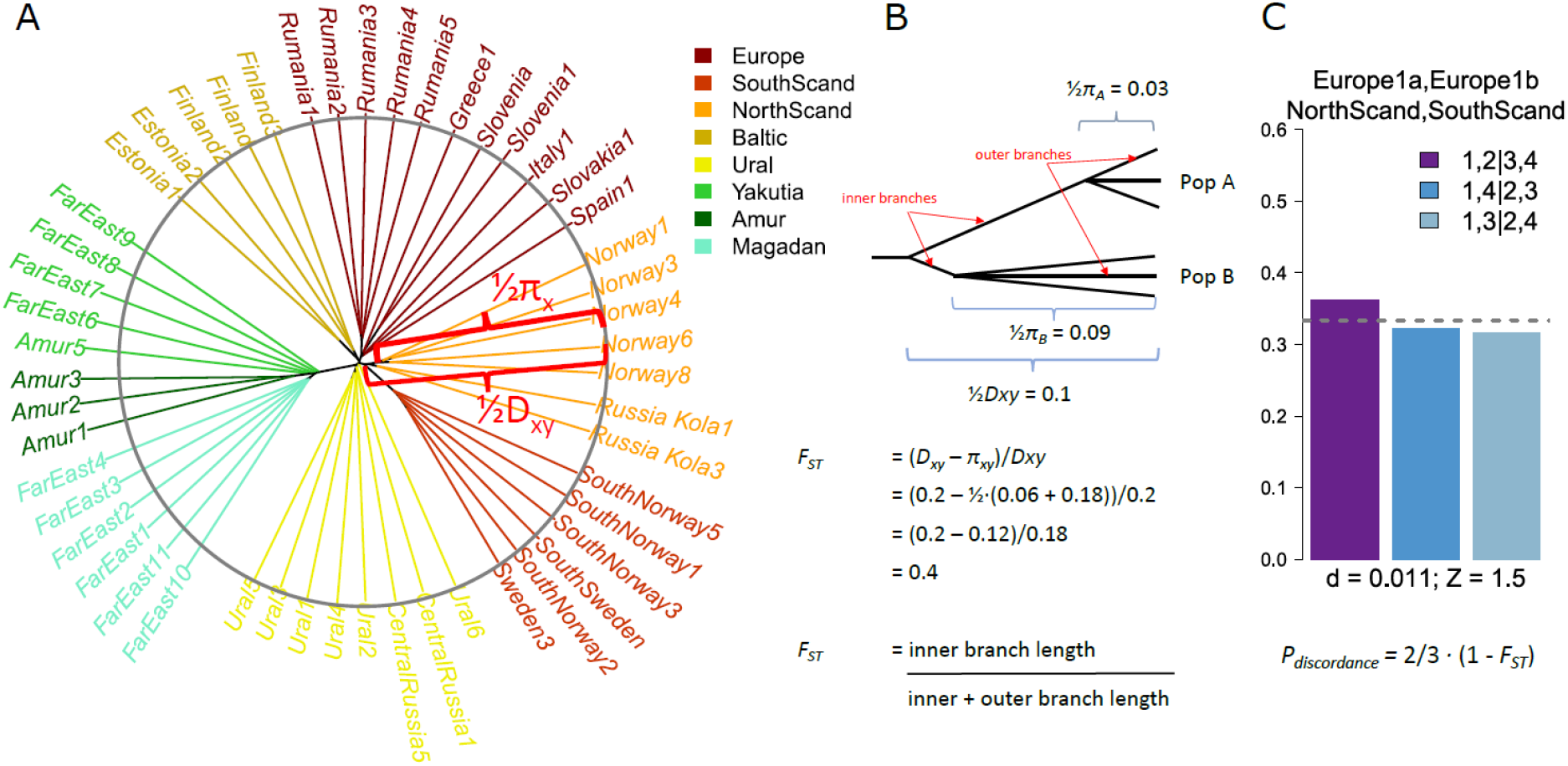
Inferring *F*_*ST*_ and gene tree discordance from multi-locus trees. *A*. A bioNJ multi-locus tree depicting whole-genome sequence dissimilarity between Eurasian brown bears, *U. arctos* (De Jong et al. 2023). *D*_*xy*_ denotes between-population distances, while nucleotide diversity, *π*, indicates genetic distances within populations. *D*_*xy*_ is highlighted between the populations of southern and northern Scandinavia, while *π*_*x*_ highlighted for the population of northern Scandinavia only. The circle has been added to aid visual interpretation, and serves as ‘isocline’, highlighting that multi-locus sequence dissimilarity estimates are not affected by genetic drift and hence translate into ultrametric trees. Eastern Eurasian bears (‘Yakutia’, ‘Amur’ and ‘Magadan’) separated from western Eurasian bears in the more distant past, and hence stand out due to the accumulation of novel mutations. ***B***. Schematic visualisation of inferring *F*_*ST*_ from proportional branch lengths. **C**. In case of a recent population split, when mutations are negligible, the amount of gene tree discordance can be estimated from *F*_*ST*_ using the formula: 2/3·(1-*F*_*ST*_). Depicted are the frequencies of the three quartet tree topologies for four brown bear populations in Europe and Scandinavia, indicating that the vast majority (approximately 64%) of the quartet tree topologies are discordant (i.e., not consistent with the species tree in A). Population ‘Europe 1a’ is represented by individual ‘Spain1’.

There are two main approaches to calculate *D*_*xy*_ and *π*_*xy*_. One approach, suitable for genomic datasets with low sample sizes, is to calculate sequence dissimilarity among all possible pairs of individuals, and subsequently infer mean values for between-population and within-population comparisons. For larger datasets, with many individuals per population, *π*_*xy*_ and *D*_*xy*_ can also be derived from allele frequency data. Given *L* biallelic sites, with minor allele frequencies *p*_*x*_ and p_y_ in populations X and Y, estimates of *π*_*xy*_ and *D*_*xy*_ are given by (Box 1):

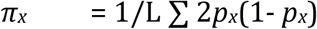

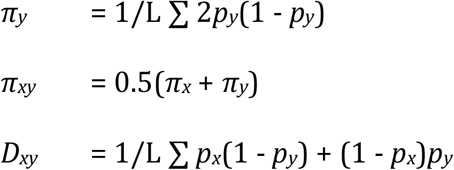

As a numerical example for a single locus, consider that two haplotypes randomly drawn from population *X* differ on average 6 out of 100 sites; that for any two haplotypes drawn from population *Y*, this number is 18 out of 100 sites; and that for any haplotype from population *X* compared to any haplotype of population *Y* this number is on average 20 differences out of 100 sites in total. We can infer:

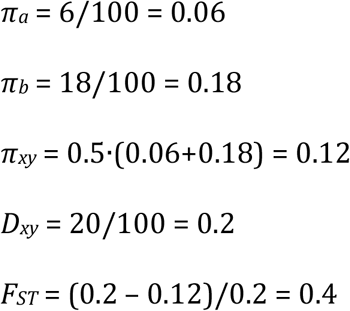

This *F*_*ST*_-value of 0.4 indicates that the pairwise sequence dissimilarity within the two populations is on average 40% smaller than the sequence dissimilarity between populations (i.e., 70% for population *X*, and 10% for population *Y*).

### Comparing Hudson *F*_***ST***_ **to other *F***_***ST***_**-metrics**

Hudson *F*_*ST*_ can be compared to other *F*_*ST*_-metrics (Box 1) by rewriting these alternative metrics as functions of *D*_*xy*_ and *π*_*xy*_. We then find that the *F*_*ST*_-metrics of Wright (1943) (Box 2), Nei (1973) (SI 1) and Nei (1977) (SI 2) differ with regard to the denominator, but have the same numerator as Hudson *F*_*ST*_, which equals Nei’s *D* (Nei 1972) (SI 3):

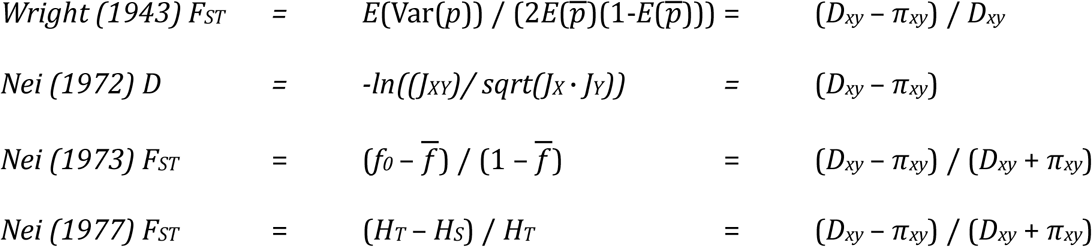

As indicated by their shared numerator, all *F*_*ST*_-metrics measure net divergence (*D*_*a*_ = *D*_*xy*_ -*π*_*xy*_). When distances are calculated from an unbiased selection of monomorphic and polymorphic sites (i.e., not from SNP datasets), *D*_*a*_ is equivalent to Nei’s *D* (Nei 1972; Nei 1973; Chakraborty and Nei 1977) (SI 3, Fig. S1).

In the classic definition of Wright (1943), net divergence is expressed as the observed variance in allele frequencies between two populations, also known as the *f*^*2*^-statistic (Reich et al. 2009; Peter 2016):

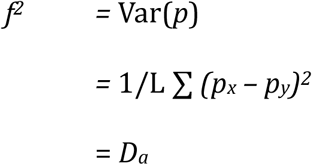

This *f*^*2*^-statistic has useful linear properties, which permits drift calculations (Fig. 2, SI 4) (Felsenstein 1981; Patterson et al. 2012).

**Figure 2.**
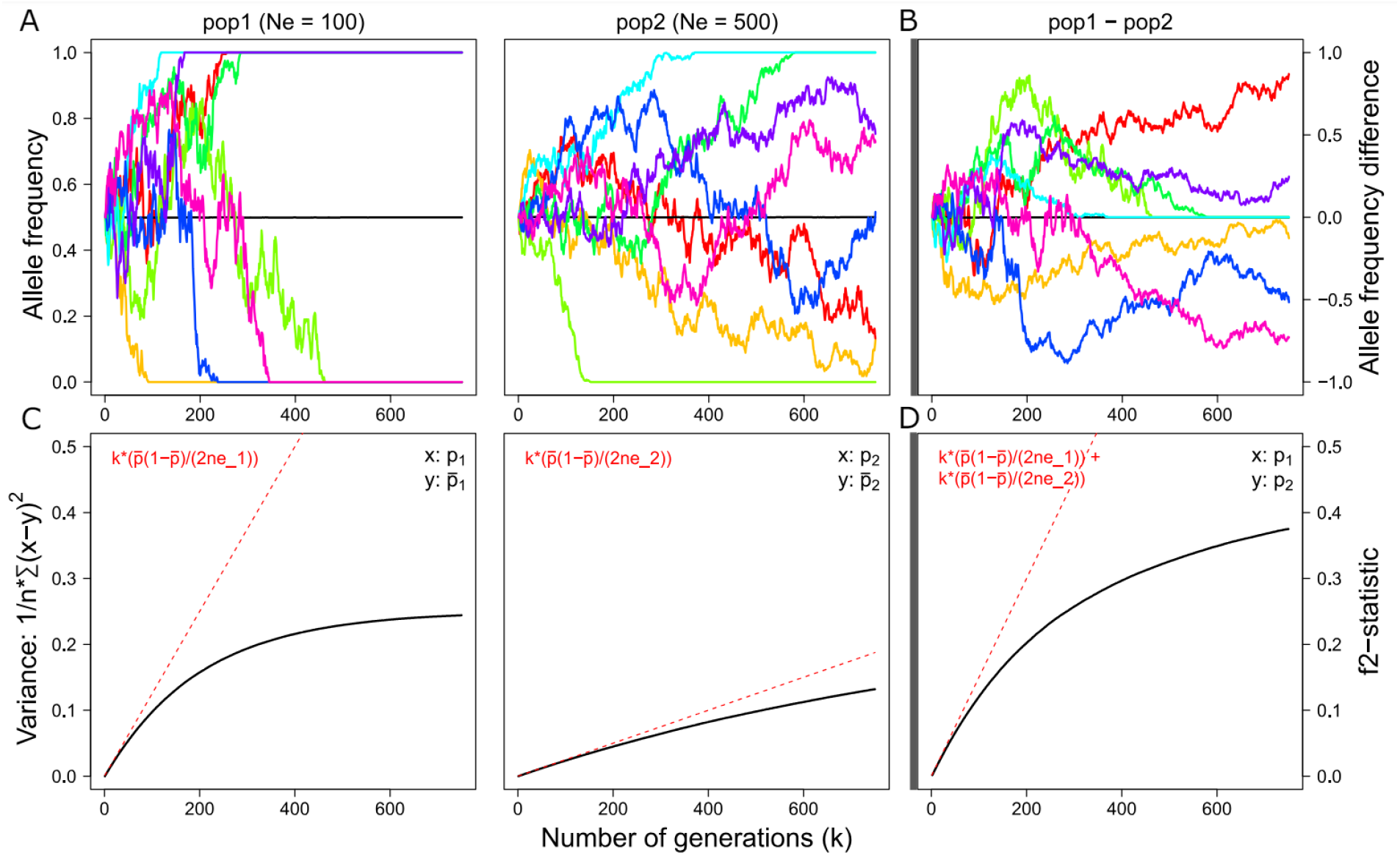
Allele frequency differences (Δ*p*) through time. **A**. Simulated allele frequency fluctuations through time for biallelic sites with an initial minor allele frequency of 0.5, given a *N*_*e*_ of 100 (left) and 500 (right) diploid individuals. The black line depicts the mean allele frequency, 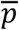, averaged over n = 100,000 sites. Note that 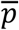 remains equal to the initial minor allele frequency. The coloured lines represent seven randomly chosen sites (same for B and C). **B**. Simulated Δ*p* between the two populations for biallelic sites. The black line depicts the mean Δ*p* (i.e., *Δp* = *p*_*1*_ – *p*_*2*_) averaged over n = 100,000 sites. Note that *Δp* remains on average zero. **C**. Simulated variance in allele frequencies (i.e., 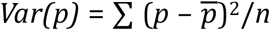 through time within the two populations, averaged over 100,000 sites. The red line indicates the expected variance in case allele frequencies were not bounded by 0 and 1. **D**. Simulated variance in Δ*p* (i.e., *f*^*2*^ = (∑ (*p*_*1*_ – p_2_)^2^)/n) through time <u>between</u> the two populations, averaged over n = 100,000 sites. Note that *f*^*2*^ is the sum of *Var(p*_*1*_) and *Var(p*_*2*_). Not depicted: Wright’s *F*_*ST*_, which is consistent with Hudson’s *F*_*ST*_, normalises the *f*^*2*^-statistic by dividing it by the maximum attainable value (i.e.,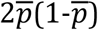).

The incongruence of the denominators indicates that the difference between the various *F*_*ST*_-metrics arises during the normalisation step, when *D*_*a*_ is converted into a proportional estimate such that it ranges between 0 and 1. This incongruence reflects a fundamental question. *F*_*ST*_ indicates a reduction in heterozygosity in subpopulations (*S*) relative to the total population (*T*), but how should the total population be defined? (Weir 2012).

Most *F*_*ST*_-metrices include within-population distances (*π*_*xy*_) when estimating distances in the total population. In contrast, and as indicated by the absence of *π*_*xy*_ in its denominator, Hudson *F*_*ST*_ does not consider within-population distances when estimating distances in the total population. This returns an intuitive proportion, namely the reduction of within-population distances relative to between-population distances.

When inferred from a sufficient number of independent loci, such that single-locus stochastics are cancelled out, *D*_*xy*_ is not affected by genetic drift (Box 3, Fig. 3A), but instead depends only on the nucleotide diversity of the ancestral population (*π*_*anc*_) and subsequent novel mutations. As a consequence, dividing *D*_*a*_ by *D*_*xy*_ returns a biologically meaningful ratio. The ratio denotes either (in case of recent divergence) the proportion of *π*_*anc*_ that has been lost by genetic drift or alternatively (in case of constant population size) the proportion of *D*_*xy*_ that can be attributed to novel mutations rather than to ancestral variation (see also section ‘Demographic inference using Hudson *F*_*ST*_’).

**Figure 3.**
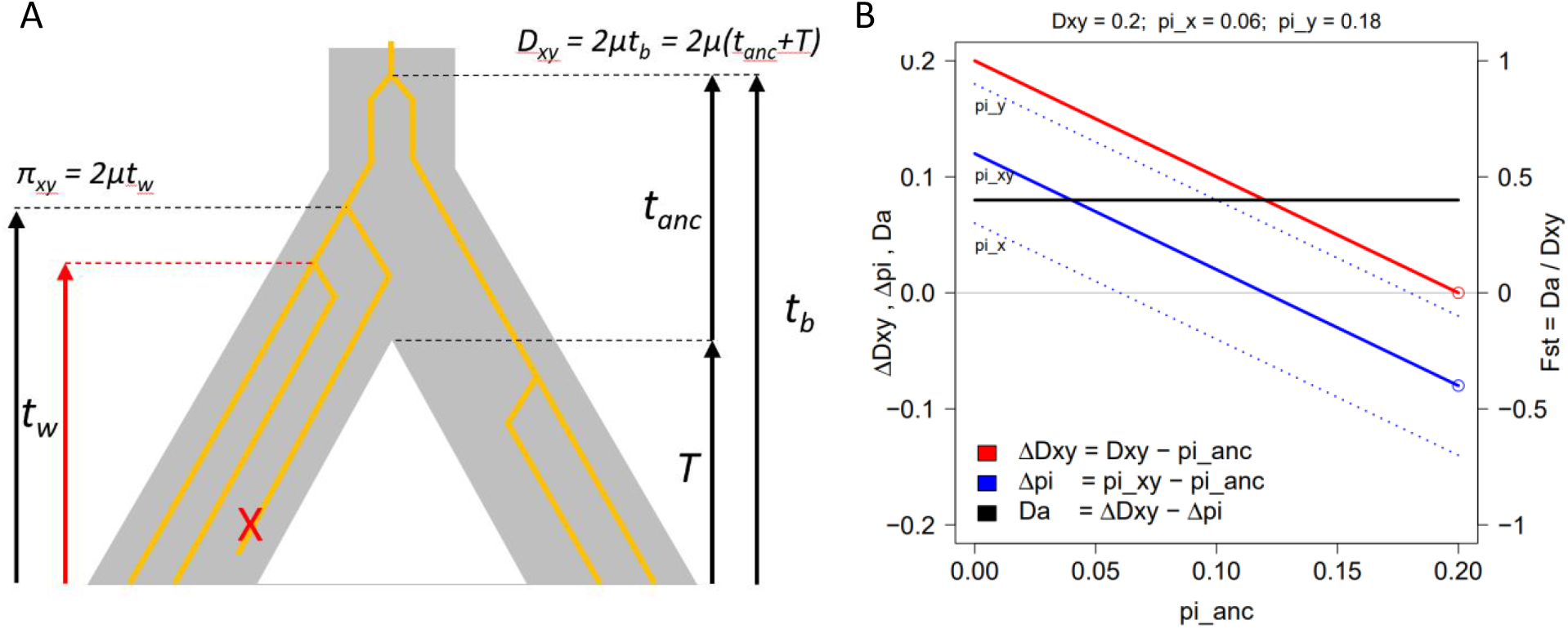
*F*_*ST*_ measures the difference in increase of *D*_*xy*_ and *π*_*xy*_ since a population split. **A**. This simplified diagram, with grey representing the species tree and orange representing a single gene tree (or ‘c-locus’ tree) depicts how the loss of variation through genetic drift can lower coalescence times within populations (*t*_*w*_) while leaving coalescence times between populations (*t*_*b*_) unaffected. It can indeed be shown that, when averaged over many c-loci, *t*_*b*_ and *D*_*xy*_ are not affected by genetic drift (Box 3). **B**. Line graph depicting the range of underlying demographic scenarios given two populations, *X* and *Y*, with a nucleotide diversity (π, or ‘pi’) of 0.06 and 0.18, and an absolute genetic distance (*D*_*xy*_) of 0.2. Depending on the nucleotide diversity of the ancestral population (*π*_*anc*_ or ‘pi_anc’), *D*_*xy*_ and *π*_*xy*_ (mean of π_x_ and π_y_) either increased and/or decreased from their starting point, *π*_*anc*_, to reach their current values. Net divergence (*D*_*a*_, black line) denotes the difference between Δ*D*_*xy*_ (red line) and Δ*π*_*xy*_ (solid blue line). *F*_*ST*_ normalises *D*_*a*_, such that it ranges between 0 and 1, by dividing it by *D*_*xy*_. Indicated by the circles on the right is one particular scenario in which the population split occurred so recently that novel mutations are absent (*π*_*anc*_ = 0.2, Δ*D*_*xy*_ = 0 and Δ*π*_*xy*_ = -0.08).

### Hudson *F*_***ST***_ is robust to sample size variation

Another asset of the Hudson *F*_*ST*_-estimator is that it is robust to sample size variation. By defining *π*_*xy*_ as the average of population averages (i.e., 0.5(*π*_*x*_ + *π*_*x*_)), the metric sidesteps potential biases which could arise from differences in sample number across populations (Bhatia et al. 2013).

Moreover, Hudson *F*_*ST*_-estimates can, in theory, be inferred with sample sizes as low as one individual per population. This is possible because genome-wide estimates of sequence dissimilarity are expected to be ultrametric. These expectations rely on the assumptions of neutrality, mutation rate constancy and absence of gene flow. For whole-genome comparisons between closely related species (or populations), the first two assumptions are realistic, such that deviations from ultrametricity can be attributed to gene flow (Cavalli-Sforza and Piazza 1975).

Within populations, *π*-estimates are expected to equal genome-wide heterozygosity (*He*), but only in the absence of cryptic population structure (i.e., panmixia) and only in the absence of close-kin matings (i.e., *F*_*IS*_ = 0). To validate these assumptions, it is advisable to include at least two or three individuals per population (Willing et al. 2012). In addition, it should be noted that ultrametricity is only observed when distances are based on a sufficient number of independent loci, such that single-locus stochastics are cancelled out.

In contrast to Hudson *F*_*ST*_, the *F*_*ST*_-metrices of Wright (1943), Nei (1973), and Nei (1977) can be sensitive to sample size variation, depending on how their components 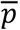, *H*_*T*_ and 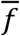 are derived (Willing et al. 2012). Biases arise if these parameters are obtained by averaging over all samples instead of averaging over population averages (Bhatia et al. 2013). In the particular case of the *F*_*ST*_-estimator of Wright (1943), bias is introduced when the maximum attainable variance (the denominator) is estimated from per-locus allele frequencies, rather than from the mean frequency across all loci (Box 1, 2).

As a numerical example, consider a single biallelic locus with minor allele frequencies, *p*_*A*_ and *p*_*B*_, of 0.1 and 0.2 in populations A and B, respectively. Because the two populations have different levels of genetic diversity, a sample set of fifteen *A*-individuals and five *B-*individuals will return a different 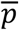 -estimate (i.e., (15·0.1+5·0.2)/20 = 0.125) than a sample set of five *A*- and fifteen *B*-individuals (i.e., (5·0.1+15·0.2)/20 = 0.175). The same applies for the parameters *H*_*T*_ and 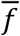, which are functions of 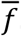. To avoid such biases, 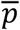, needs to be calculated as the average of the population averages: 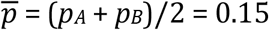 = (*p*_*A*_ + *p*_*B*_)/2 = 0.15.

### Demographic inference using Hudson *F*_***ST***_

What exactly can we infer from an *F*_*ST*_-estimate about the underlying evolutionary processes which caused the observed level of population differentiation? It is well-known that, under certain conditions, *F*_*ST*_-estimates may serve as an indirect measure of gene flow, as given by the formula: *F*_*ST*_ = 1/(*4N*_*e*_*m* + 1) (Slatkin 1993), with *N*_*e*_ and *m* denoting effective population size and migration rate, respectively. However, this formula is foremost a theoretical construct with little bearing to real-world data, as it assumes migration-drift equilibrium, which most natural populations will never reach (Whitlock and McCauley 1999).

More generally valid predictions of *F*_*ST*_ can be made by describing *F*_*ST*_ as a function of mutation rate (*μ*), effective population size (*N*_*e*_), and effective population split time (*T*_*e*_). The latter factor, *T*_*e*_, can be more recent than the actual split time (*T*) owing to subsequent gene flow (Mountain and Cavalli-Sforza 1997). Unfortunately, population-genetic theory only allows for hypothetico-deductive reasoning, namely to predict *F*_*ST*_ given a hypothetical evolutionary scenario. The opposite reasoning, where we aim to induce the hypothesis (evolutionary scenario) from the observation (*F*_*ST*_-estimate), is not possible. The reason is equifinality, with different sequences of events (i.e., different combinations of *Ne, T*_*e*_ and *μ*) being able to produce an identical outcome (i.e., same *F*_*ST*_-estimate).

The equifinality of *F*_*ST*_ becomes evident when we reformulate Hudson *F*_*ST*_ by rewriting one of its two components, *π*_*xy*_, as the sum of the nucleotide diversity of the ancestral population (*π*_*anc*_) and its subsequent increase or decrease (Δ*π*_*xy*_ = *π*_*xy*_ – *π*_*anc*_) (SI 5):

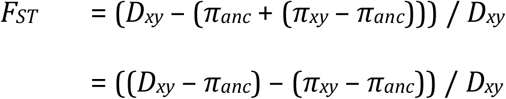

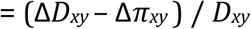

This reformulation shows that Hudson *F*_*ST*_-estimates measure the difference in increase of both *D*_*xy*_ and *π*_*xy*_ since the population split (Fig. 3B). However, these two components, Δ*D*_*xy*_ and Δ*π*_*xy*_, cannot be inferred separately. There are an infinite number of potential underlying demographic scenarios (i.e., various combinations of Δ*D*_*xy*_ and Δ*π*_*xy*_), all of which can explain the observed level of population differentiation equally well (Fig. 3B).

For instance, for our numerical example (see section ‘Calculating Hudson *F*_*ST*_’), the *F*_*ST*_-value of 0.4 indicates that *π*_*xy*_ increased 40% less than *D*_*xy*_ since the population split (Fig. 3B). One potential underlying scenario is that the population event occurred so recently that novel mutations are absent, such that *D*_*xy*_ remained equal (i.e., *ΔDxy* = 0) (Fig. 3B). According to this scenario, the ancestral population had a nucleotide diversity of 0.2, of which 70% and 10% (thus, 40% on average) was lost by populations *X* and *Y*, respectively (Fig. 3B). Because in this scenario *T* is practically zero (i.e., very recent), there must have been a severe reduction in *N*_*e*_, especially in population *Y*.

Another potential underlying scenario, on the opposite extreme, is that the ancestral population was devoid of genetic variation (i.e., *π*_*anc*_ = 0), such that the observed sequence dissimilarities are caused entirely by novel mutations and not by ancestral variation. According to this scenario, and assuming a mutation rate (*μ*) of 10^−6^, the population split would have occurred 100,000 generations ago, which is the time needed for mutations to increase pairwise sequence dissimilarity between populations from 0 to 0.2. In nature, *μ* and *π* are typically two orders of magnitudes lower, but the final split estimate would be the same.

A third scenario, among the many more, is that nucleotide diversity remained equal on average (i.e., Δ*π*_*xy*_ = 0). In this scenario, the ancestral population had a nucleotide diversity of 0.12, after which *π*_*x*_ increased to 0.18 while *π*_*y*_ decreased to 0.06 (Fig. 3B).

The shared feature of any of the possible underlying scenarios is that at the time of the split event, at *T*_*e*_ generations ago, *D*_*xy*_ and *π*_*xy*_ both equal the nucleotide diversity of the ancestral population (*π*_*anc*_). This level starting point, *π*_*anc*_, is a product of two factors: mutation rate *(μ*), and the mean coalescence time in the ancestral population (*t*_*anc*_):

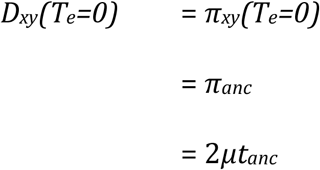

After the population split, the mean coalescent time of a pair of haplotypes from different populations (*t*_*b*_) increases linearly with split time (*T*_*e*_). After all, if the mean coalescence time in the ancestral population (*t*_*anc*_) is, say, 4000 generations, and if two subpopulations subsequently become separated for, say, 2000 generations, then the mean coalescence time between the two populations (*t*_*b*_) will simply increase to 6000 generations. Therefore, assuming mutation rate constancy, we can infer that *D*_*xy*_ is proportional to the sum of *T*_*e*_ and *t*_*anc*_ (Wakeley 2000; Hancock and Blackmon 2020):

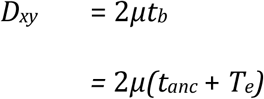

We can furthermore infer that *D*_*xy*_ depends only on the population size of the ancestral population (*Ne*_*anc*_), not on population size changes after the population split (Box 3):

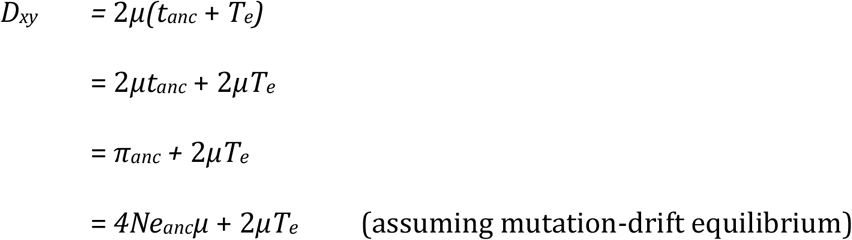

The reason that *D*_*xy*_ is not affected by population size changes after the population split, is that the loss of alleles owing to genetic drift reduces the mean coalescent times within populations (*t*_*w*_) by a certain factor (*ω*), but it does not alter the mean coalescent time between populations (*t*_*b*_) (Fig. 3A, Box 3):

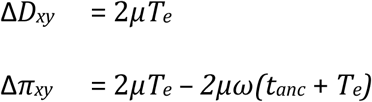

Since the factor 2*μT*_*e*_, which represents novel mutations, occurs in Δ*D*_*xy*_ as well as Δ*π*_*xy*_, it is cancelled out from the *F*_*ST*_-estimate, indicating that *F*_*ST*_ denotes the reduction of coalescence time within populations, irrespective of the split time (*T*_*e*_) (Fig. 3A):

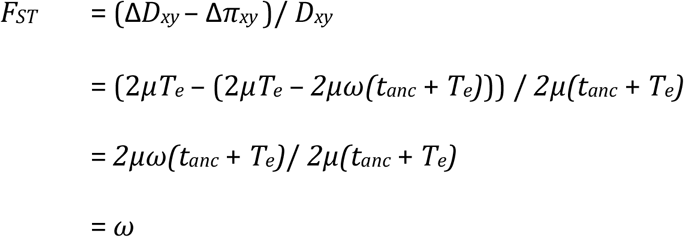

This is what we should expect to find, because novel mutations following the population split increase sequence dissimilarity within populations (*D*_*xy*_) at the same rate as they increase sequence dissimilarity within populations (*π*_*xy*_). For instance, consider a split event which separates two haploid, asexual populations with non-overlapping generations. If all these individuals consistently produce one offspring each generation by selfing, such that population sizes remain equal whilst simultaneously avoiding genetic drift (i.e., no vicariance events), at any point in time two sequences within a population are expected to differ as much as two sequences from either population. This illustrates that the difference between *D*_*xy*_ and *π*_*xy*_ arises *solely* due to the loss of genetic variation though genetic drift, not due to mutations.

To return to our numerical example, if we assume a mutation rate (*μ*) of 10^−6^ per generation, we can infer that the mean coalescence time within populations is 60,000 generations (i.e., *t*_*w*_ = 0.12/(2·10^−6^)), which is 40% lower than the mean coalescence time between populations (*t*_*b*_ = 0.2/(2·10^−6^) = 100,000 generations). Without specifying the exact underlying demographic scenario, the *F*_*ST*_-value of 0.4 indicates that *t*_*w*_ has been reduced by 40% relative to *t*_*b*_, owing to genetic drift following the population split (Slatkin 1991).

### The special case of *ΔDxy* = 0

Among the countless scenarios which can produce the same *F*_*ST*_-estimate, one particular scenario entails that the split event occurred so recently that novel mutations are negligible. Because mutation rates are generally low, intraspecies population divergence can often be approximated by this theoretical scenario, especially for bottlenecked populations, when loss of genetic variation greatly outweighs novel mutations (Cavalli-Sforza and Edwards 1967).

In case of such a very recent population split, such that *ΔDxy* ≈ 0, and assuming equal population sizes (i.e., *N*_*e_x*_ = *N*_*e_y*_), the predictions for *D*_*xy*_ and *π*_*xy*_ simplify to:

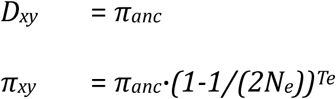

Substituting these factors, *F*_*ST*_ can be expressed as a function of *T*_*e*_ and *N*_*e*_, denoting the average proportion of ancestral genetic variation (*π*_*anc*_) that after the split event has been lost in either population as a result of genetic drift:

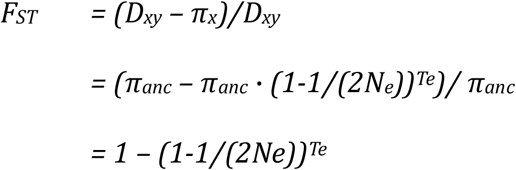

Since it is usually impossible to attribute the loss of genetic variation to a specific combination of *N*_*e*_ and *T*_*e*_, the best we can do is to express population split times in multiples of their haploid effective population size, also known as coalescent units (*τ*) (Degnan and Rosenberg 2009):

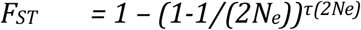

Inverting the above formula, and replacing 2*N*_*e*_ with a random high number (e.g., 1000), we can infer *τ* from *F*_*ST*_, as follows (Fig. 4):

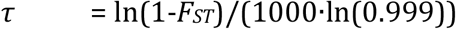

**Figure 4.**
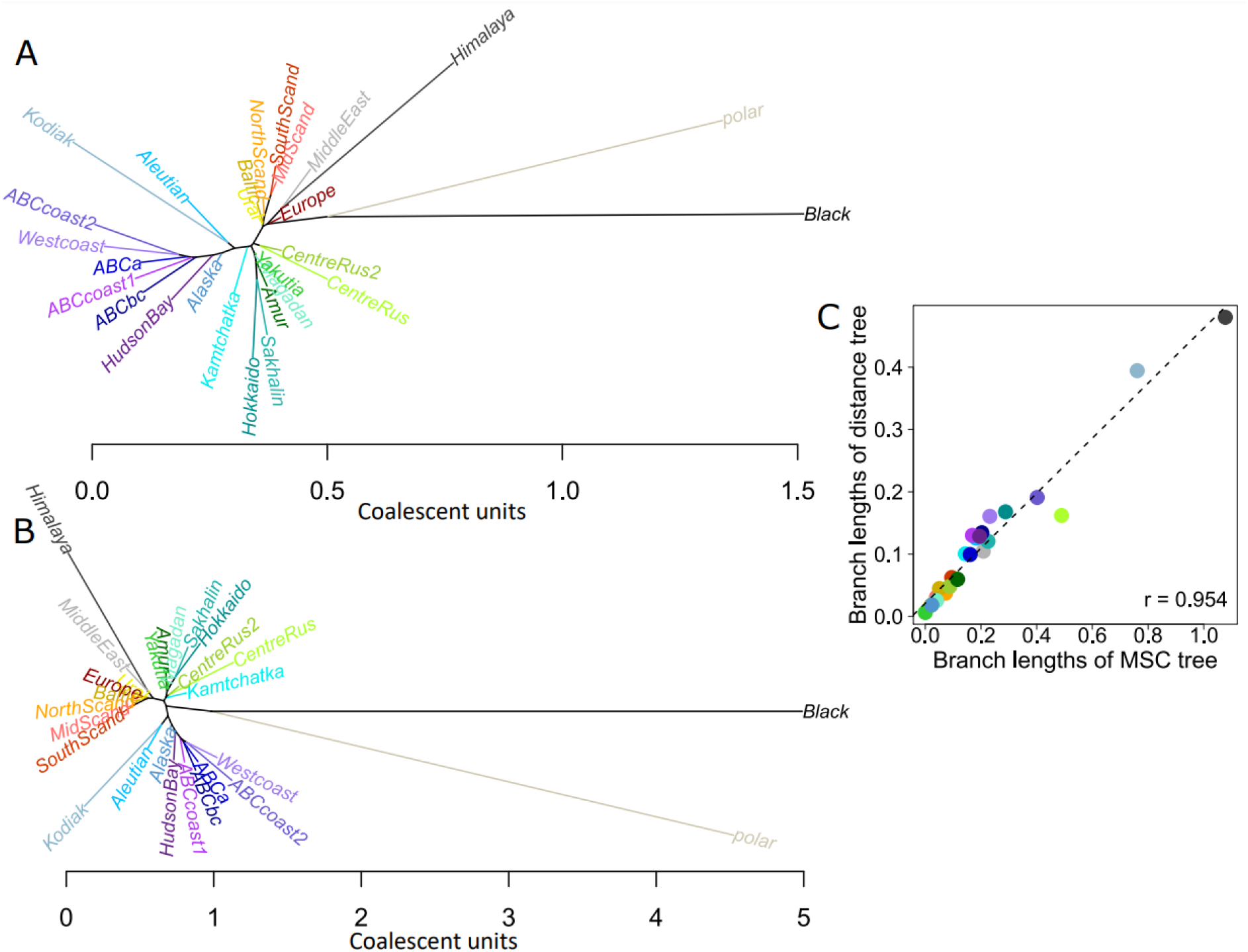
Comparison of *F*_*ST*_-based *τ*-estimates and MSC-based *τ*-estimates. A. Distance-based bioNJ-phylogeny inferred from a population-pairwise distance matrix with *τ*-estimates, inferred from Hudson *F*_*ST*_-estimates using the formula: *τ* = (ln(1-*F*_*ST*_)/ln(0.999))/1000, for a real-word dataset of brown bears (De Jong et al. 2023). **B**. Multispecies-coalescent based (MSC) phylogeny inferred by Astral (Mirarab et al. 2014) for the same dataset of brown bear individuals, calculated using an input dataset of over 3000 c-loci trees (De Jong et al. 2023). **C**. Scatterplot depicting the correlation between terminal branch lengths of the distance-based tree (see A) and the MSC-based tree (see B). While considerable inconsistency consists in terms of the exact values (possible due to data artefacts related to a bias selection of c-loci), the values are highly correlated.

For instance, an *F*_*ST*_-value of 0.1 indicates a split time of *τ* = 0.105. For 2*N*_*e*_ = 200 and 2*N*_*e*_ = 20000, this translates to 21 and 2100 generations, respectively. We would need additional information, such as timing of geological events underlying the population split, as well as an accurate estimate of long-term *N*_*e*_, to resolve which of these two scenarios, or any of the other endless possible combinations of *N*_*e*_ and *T*_*e*_, represents the true scenario (Degnan and Rosenberg 2009).

While the derivation of coalescent units in itself does little to narrow down population split times, they are useful in another respect: they allow to quantify the amount of gene tree discordance within genomes (i.e., where phylogenies from individual c-loci differ from the species phylogeny). Coalescent theory predicts that the probability of deep coalescence *P(t > T*_*e*_), when coalescent events are older than the population split, is a function of *N*_*e*_ and *T*_*e*_, and thus of *τ* (Allman et al. 2011). Each generation, the probability that two haplotypes coalesce is the inverse of the size of the haploid gene pool (2*N*_*e*_ in diploid populations), and hence the probability that two haplotypes do *not* coalesce during T generations, is given by:

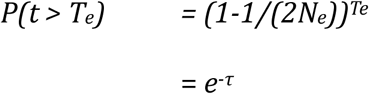

Note the similarity between this formula and the formula for *F*_*ST*_ when assuming the absence of novel mutations, i.e.; *F*_*ST*_ *= 1 – (1-1/(2N*_*e*_))^*Te*^. Comparing these two formulas, we arrive at the conclusion that, in the absence of novel mutations, *F*_*ST*_-values estimate the inverse of deep coalescence: namely, the proportion of c-loci haplotypes that *do* coalesce more recently than the population split:

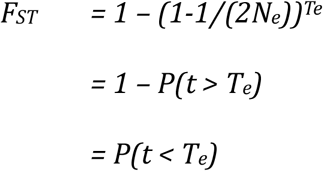

In case of deep coalescence, when coalescence events are older than the population split (i.e., *P(t > T*_*e*_)), a locus will randomly, and hence equally, represent the three unrooted quartet trees (AB|CD), (AC|BD), and (AD|BC) (Allman et al. 2011). Only one of this three quartet trees corresponds with the species tree. For the other two quartet trees, the coalescence time of two haplotypes drawn from within the same population (*t*_*w*_) will be greater than that of two haplotypes drawn from different populations (*t*_*b*_), and thus discordant (Maddison 1997; Carling and Brumfield 2008; Allman et al. 2011). Hence, we can infer that the probability of gene tree discordance (t_w_ > t_b_) as a result of incomplete lineage sorting, is given by:

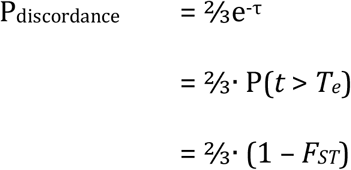

As a numerical example, consider two populations, *A* and *B*, with a pairwise *F*_*ST*_-value of 0.1, implying a split time (*τ*) of 0.105·2*N*_*e*_ generations relative to the ancestral population (see start of this section). Let’s furthermore assume that in our dataset the two populations are each represented by a single diploid individual, one carrying haplotypes A1 and A2, the other haplotypes B1 and B2. From the split time estimate, we can infer that for (e^-0.105^) 90% of c-loci, the two haplotypes in an individual’s diploid genome have split times predating the population split. In other words, for 9 out of 10 c-loci, haplotypes A1 and A2 (and similar B1 and B2) coalesce further back in time than the population split. Note that this proportion can also be inferred directly from the *F*_*ST*_-value:

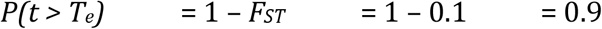

We can furthermore estimate that two-thirds of deep coalescing c-loci (i.e., 2/3 · 90% = 60%) will be discordant, such that haplotype A1 coalesces with haplotype B1 or B2 prior to coalescing with haplotype A2. The remaining 40% of c-loci, a slight majority among the three types of quartet trees, will support the quartet topology corresponding to the species tree topology, such that haplotype A1 coalesces first with haplotype A2. For reference: prior to the population split, this proportion was 1/3th (33.3%); given sufficient time, and assuming absence of gene flow, this proportion will eventually converge to 1, such that all loci support the topology of the species tree.

An F_ST_-value of 0.1 is within the normal range of *F*_*ST*_-values typically observed for intraspecific divergence. As an arbitrary example from real-world data, the genetic differentiation of Scandinavian and European brown bears (*U. arctos*) returns an *F*_*ST*_-value of close to 0.1, as can also be inferred from the proportional length of inner branches (Fig. 1A). While the multi-locus tree reveals distinct clusters which clearly separate European individuals from Scandinavian individuals, indicating reciprocal monophyly (Fig. 1A), each individual c-locus tree (i.e., gene tree) is expected to be highly polyphyletic, with close to 65% of the quartet trees being discordant (Fig. 1C). For comparison, the insular population of Kodiak bears experienced a severe population bottleneck, such that the *F*_*ST*_-value relative to their closest relatives on the nearby mainland equals 0.4 (de Jong et al. 2023). Even for this pair of highly diverged populations, considerable polyphyly is expected within c-locus trees, with around 40% (i.e., 2/3·(1-0.4)) of quartet trees being discordant.

It should be stressed that above considerations on the relationship between *F*_*ST*_ and *τ* (and therewith the extent of gene tree discordance) only apply when novel mutations are negligible (i.e., *ΔD*_*xy*_ ≈ 0) (Fig. 5). The derivations lead to underestimates of coalescence times when applied to population pairs for which novel mutations cannot be ignored (Fig. 5). This may be because of deep split times (high *T*_*e*_) and/or because of low nucleotide diversity in the ancestral population (low *π*_*anc*_) (Fig. S2). In either case a relatively high proportion of *D*_*xy*_ can be attributed to novel mutations, which violates the assumption that novel mutations are negligible.

**Figure 5.**
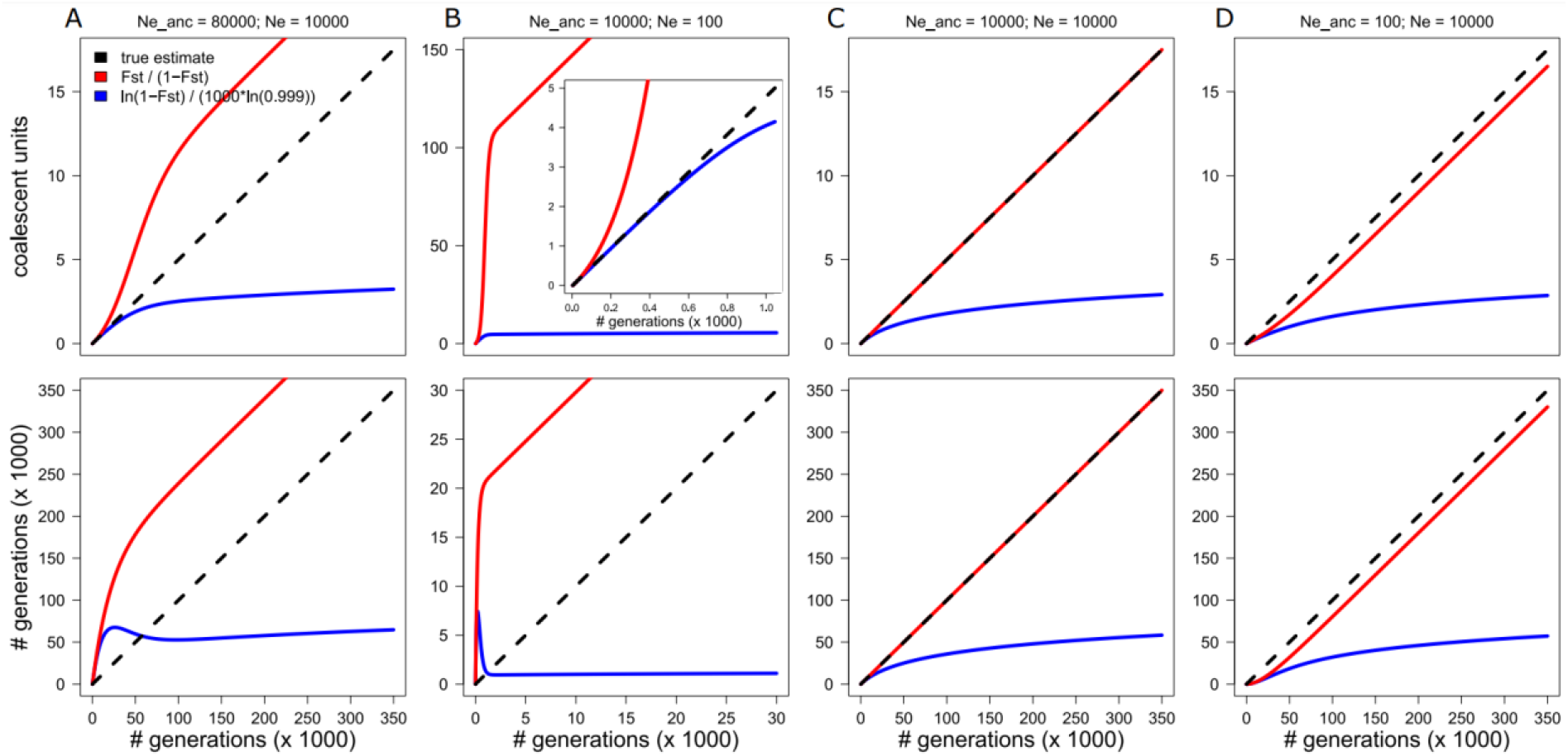
No universal relationship exists between *F*_*ST*_ and *τ* (coalescent units). Line graphs depicting split times estimates in coalescent units (*τ*, top row) and number of generations (*T*, bottom row), inferred from *F*_*ST*_-estimates for four different evolutionary scenarios. The true estimates (black dashed line) are based on an iterative function (see SI 7), assuming a per-generation mutation rate of 10^−8^. *τ* has been inferred from *F*^*ST*^ either assuming population size constancy (*τ*^*1*^, red solid line) or alternatively assuming absence of novel mutations (*τ*^*2*^, blue solid line). Subsequently, *T* was inferred from *τ* using the formula: *T* = *τ*·*2N*^*e*^, with *N*^*e*^ = *π*/(4*μ*). **A**. and **B**. If *N*^*e*^ decreases after a population split, *τ*^*2*^ is accurate for low *τ*^*2*^. However, because the extant populations are not yet in mutation-drift equilibrium, *π*/(4*μ*) is an overestimate of true *N*^*e*^, and hence so is *T*. Hence, *N*^*e*^-estimates (and therewith *T*-estimates) should not be obtained assuming mutation-drift equilibrium. Inset zooms in on the first 1000 generations. **C**. If *N*^*e*^ remains constant, and assuming mutation-drift equilibrium at the time of the split event, *τ*^*1*^ accurately estimates the true number of coalescent units, regardless of split time. **D**. If *N*^*e*^ increases after a population split, *τ*^*1*^ and *τ*^*2*^ are underestimates.

### The special case of Δ*π*_***xy***_ = 0

For comparison, let us consider an alternative scenario, namely the scenario in which nucleotide diversity remains on average constant through time (i.e., Δ*π*_*xy*_ = 0). For this scenario, in which net divergence (*D*_*a*_) is determined by *ΔD*_*xy*_ only and thus directly proportional to split time (Nei and Li 1979; Slatkin 1991; Wakeley 2000), it is easy to see that the number of coalescent units, *τ* (i.e., split time expressed in 2*N*_*e*_), is simply given by the ratio between net divergence and nucleotide diversity:

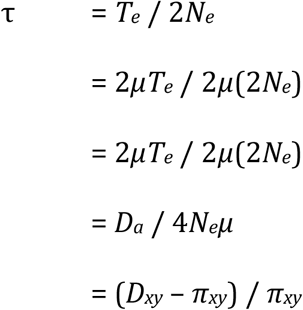

As a numerical example, consider an ancestral diploid population with *N*_*e*_ = 100.000 and *μ* = 1·10^−8^. In case of mutation-drift equilibrium, in which the loss of genetic variation through drift is balanced out by the gain of new variation through novel mutations, the expected nucleotide diversity is 0.004 (i.e., *π* = *4N*_*e*_*μ* = 0.004). After a population split event, the absolute genetic distance between the two sister populations is expected to increase to 0.005 within a time span of 50,000 generations (i.e., *D*_*XY*_ = *π* + *2Tμ* = 0.004 + 2 · 50000 · 10^−8^ = 0.005). We can infer from *D*_*xy*_ and *π*_*xy*_ that the number of coalescent units is 0.25 (i.e., *τ* = *D*_*a*_/*π* = (0.005 – 0.004)/0.004 =0.25). This is the correct estimate, as a split time of 50,000 generations is indeed a quarter of the haploid effective population size (2·100000 = 200000).

Note that this formula to infer coalescent units has the same numerator as the formula for Hudson *F*_*ST*_, but that the denominator is *π*_*xy*_ instead of *D*_*xy*_. Therefore, we can infer:

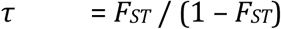

Thus, when assuming population size constancy (i.e., Δ*π*_*xy*_ = 0), we obtain a very different prediction for *τ* than when assuming absence of novel mutations (i.e., Δ*π*_*xy*_ = 0) (Fig. 5, SI 6-7). The predictions are only valid under specific circumstances, meaning that caution should be exercised when inferring *τ*-estimates from *F*_*ST*_ (Fig. 5). As an alternative approach, multispecies coalescent (MSC) based analyses can estimate *τ* from the observed gene tree discordance, without knowing the underlying demographic scenario (Fig. 4) (Degnan and Rosenberg 2009; Mirarab et al. 2021).

## CONCLUSIONS

Relative genetic distances between populations can be inferred from sequence alignment data, irrespective of sample size or sample size variation, using the formula of Hudson et al. (1992): *F*_*ST*_ = (*D*_*xy*_ – *π*_*xy*_)/*D*_*xy*_. This *F*_*ST*_-estimator measures how much smaller sequence dissimilarity within populations (*π*_*xy*_) is compared to that between populations (*D*_*xy*_). This ratio can also be read from multi-locus trees as the proportional length of inner branches. Like other *F*_*ST*_-metrics, the numerator denotes net divergence (*D*_*a*_), which is equivalent to both the *f*^*2*^-statistic and Nei’s *D* (provided realistic *D*_*xy*_ and *π*_*xy*_ estimates), and which arises after because genetic drift affects *π*_*xy*_ a population split but not *D*_*xy*_. Because different combinations of Δ*π*_*xy*_ and Δ*D*_*xy*_ can produce identical Hudson *F*_*ST*_-estimates (‘equifinality’), it is not possible to convert *F*_*ST*_ into generally valid estimate of coalescent units (*τ*). Still, if assuming absence of novel mutations (i.e., Δ*D*_*xy*_ = 0), which approximates many cases of intraspecific divergence, the expected proportion of gene tree discordance is given by: ⅔(1 – *F*_*ST*_).

### BOX 1. Calculating *F*_*ST*_ from allele frequency data

Numerical example of *F*_*ST*_-estimation given an example dataset of four biallelic loci. It is assumed that each locus had a minor allele frequency (*p*) of 0.5 when the ancestral population separated into the two extant populations (*A* and *B*):

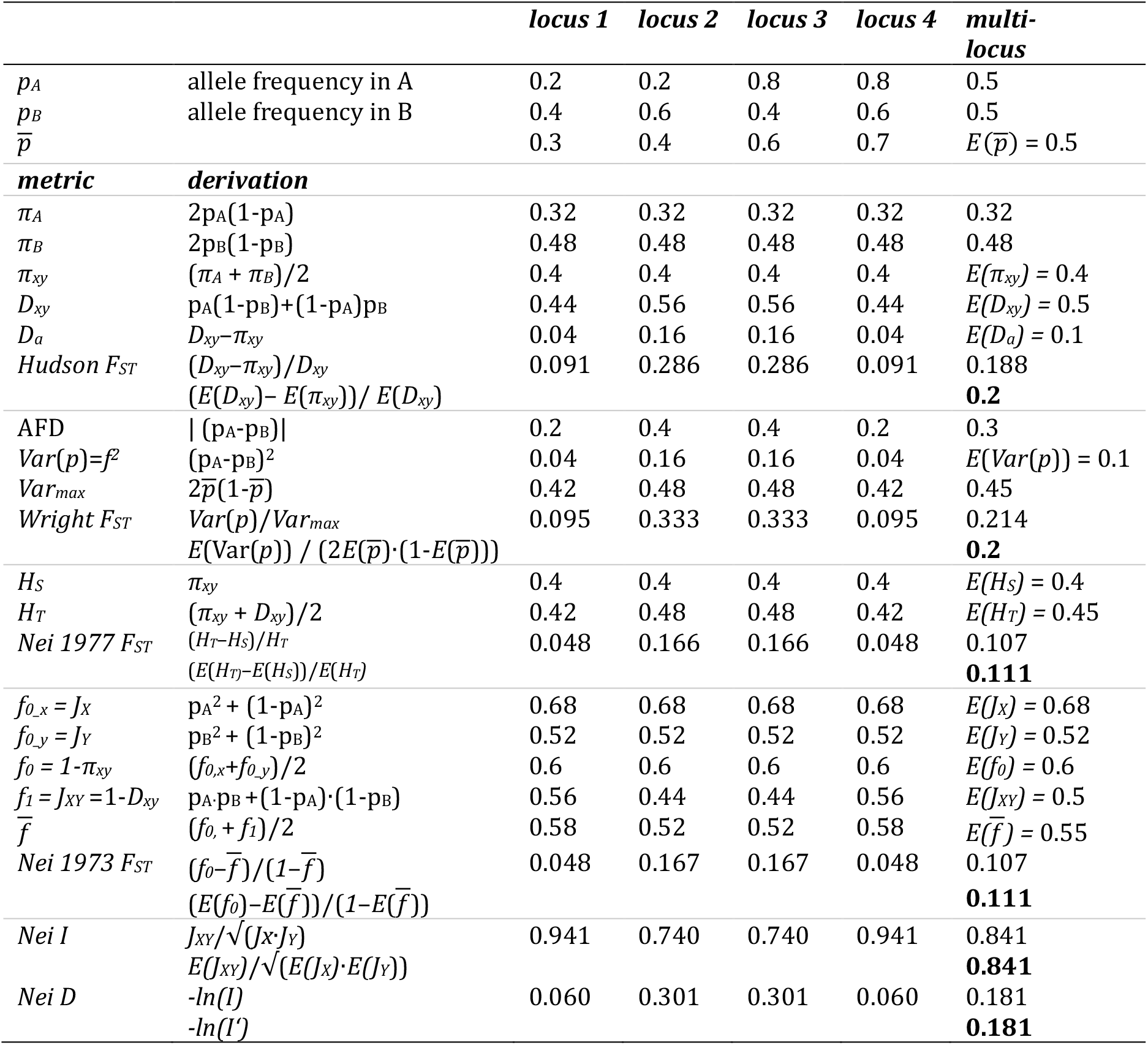

Note that *F*_*ST*_-estimates should not be obtained by averaging single-locus *F*_*ST*_-estimates, but instead need to be calculated from multi-locus parameters. Also note the discrepancy between the various *F*_*ST*_-estimators. Hudson *F*_*ST*_ and Wright *F*_*ST*_ equal 0.2, indicating that genetic distances are 20% smaller within populations than between-populations. Because *Nei’s D* is here inferred from SNP data, it overestimates *D*_*a*_.

### BOX 2.

**Comparing Hudson (1992) *F***_***ST***_ **to Wright** (1943) ***F***_***ST***_

Net divergence, *D*_*a*_, the numerator of Hudson *F*_*ST*_, can be rewritten to the *f*^*2*^-statistic, the numerator of Wright *F*_*ST*_:

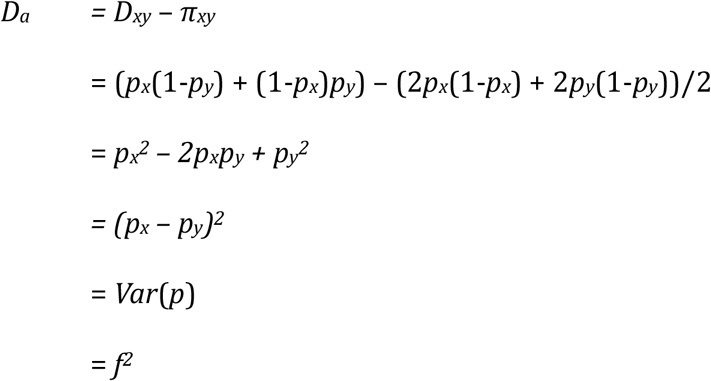

The denominator of Wright *F*_*ST*_ normalises the *f*^*2*^-statistic, by dividing the observed variance by the maximum attainable variance, when all alleles are fixed (Balloux and Lugon-Moulin 2002). For pairwise population comparisons, this upper limit is given by 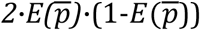 (Chen et al. 2015). Because the expected allele frequency, 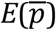, remains identical to the mean allele frequency of the ancestral population, *p*_anc_ (Fig. 2), we can infer that the denominators of Wright *F*_*ST*_ and Hudson *F*_*ST*_ are also equivalent:

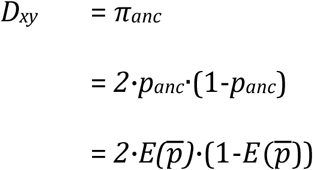

Crucially, the denominator (i.e., maximum attainable variance) need to be calculated from the mean multi-locus allele frequency, 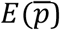, not from locus-specific allele frequencies 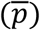 (Box 1). For sites for which allele frequencies drifted in the two populations in the same direction, the product of 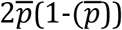 is on average biased downwards, such that the latter approach would result in an overestimate of the multi-locus *F*_*ST*_ (Box 1).

### BOX 3.

**Why are multi-locus *D*_*xy*_-estimates not affected by genetic drift?**

After a split event, genetic drift will eventually fix the ancestral variation in the two sister populations. This variation will end up in one of two states. Drift can either fix the same allele in the two populations (*D*_*xy*_ = 0), or alternatively fix different alleles in the two populations (*D*_*xy*_ = 1).

While genetic drift thus affects single-locus *D*_*xy*_-estimates, it does not affect multi-locus *D*_*xy*_-estimates, provided that a sufficient number of unlinked loci are included such that single-locus stochastics are cancelled out. The reason is that the *D*_*xy*_-values of the two states counterbalance each other. The probability of these two states is given by the probabilities of homozygosity (1-*π*_*anc*_) and heterozygosity (*π*_*anc*_) in the ancestral population, respectively. Thus, if it were for genetic drift only, *D*_*xy*_ remains equal to *π*_*anc*_:

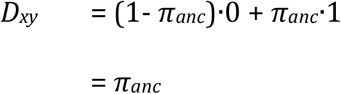

This expectation is valid irrespective of the magnitude of genetic drift in either population.

As a numerical example, consider an ancestral panmictic population which we genotyped using 1000 unlinked, biallelic SNPs with each a minor allele frequency (*p*) of 0.2. The expected heterozygosity, and thus nucleotide diversity, in this ancestral population is 2·0.8·0.2 = 0.32. Next, consider that this ancestral population splits in two sister populations, and that either population reduces in size, such that as a result of genetic drift all sites eventually become fixed for one allele. The probability that for any given site these sister populations happen to fix a different allele, is, again, 2·0.8·0.2 = 0.32.

