## Supplementary figures and information for "Calculating and interpreting *F*_*ST*_ in the genomics era"

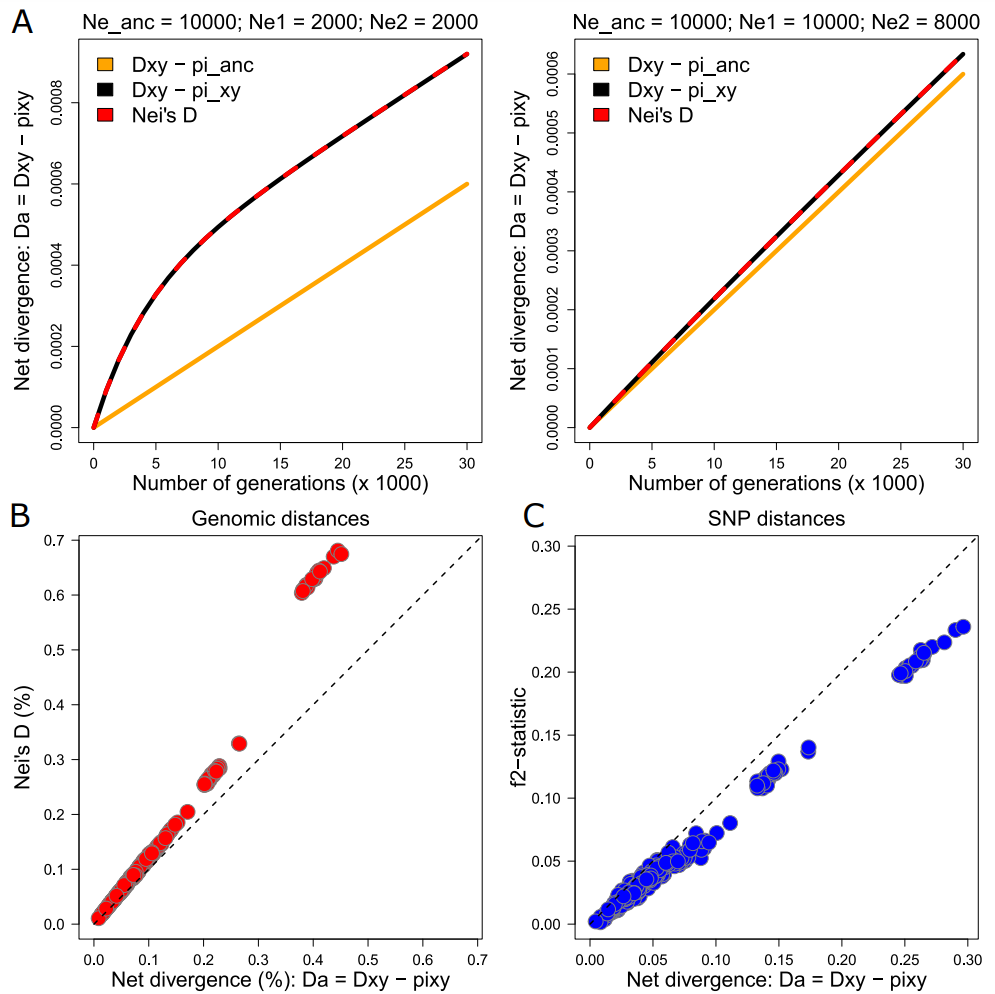

**Figure S1. Nei’s *D* and the *f^2^*-statistic are estimators of net divergence (*D_a_*). A.** Predicted values of Nei’s *D* and *D_a_* for two different scenarios. Note that Nei’s *D* equals *D_a_*. The underlying R script can be found in SI 7. **B**. Estimates of Nei’s *D* and *D_a_* inferred from sequence dissimilarity estimates for a real-world dataset of brown bears (*U. arctos*), polar bears (*U. arctos*) and American black bears (*U. americanus*) (de Jong et al. 2023). Consistent with expectations, Nei’s *D* estimates are very similar to *D_a_*-estimates, especially for intraspecific population comparisons (i.e., *D_a_* < 0.002). **C**. Estimates of *f^2^* and *D_a_* inferred from a 50K SNP dataset for the set of bear populations (de Jong et al. 2023). Consistent with expectations, *f^2^*-estimates are very similar to *D_a_*-estimates.

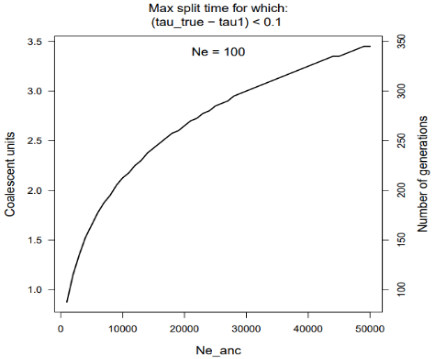

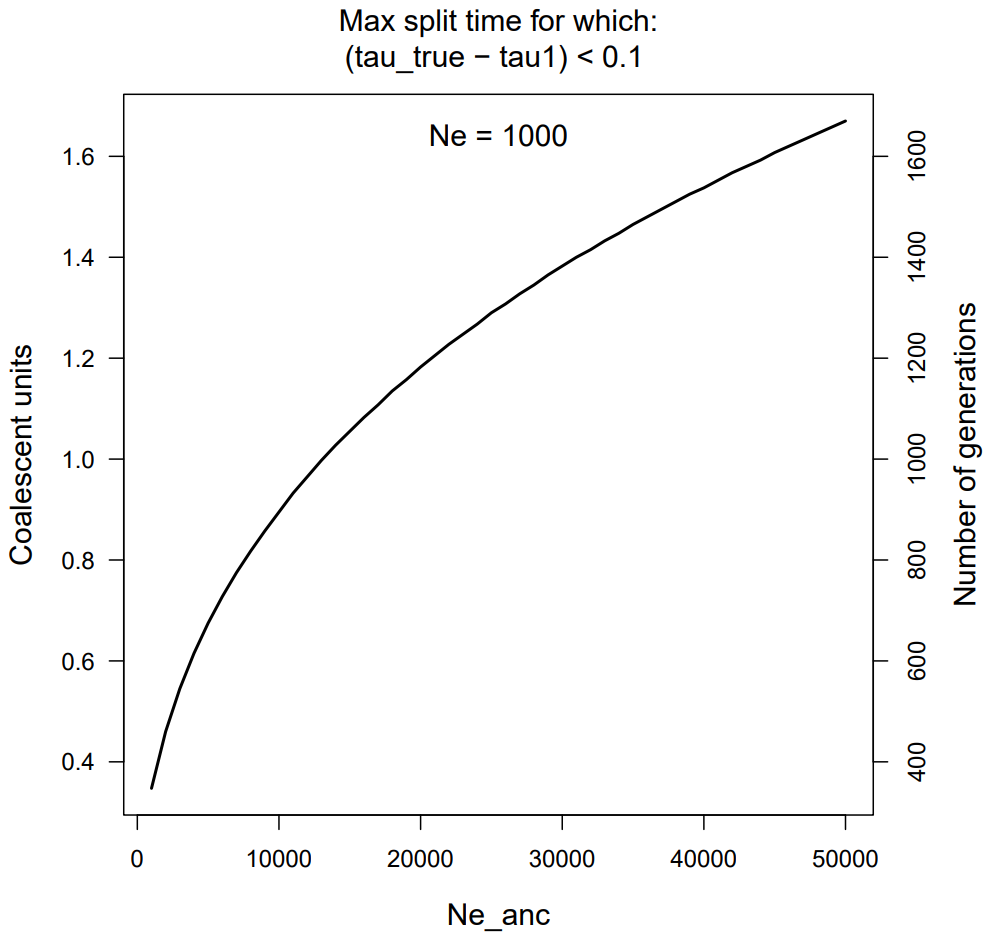

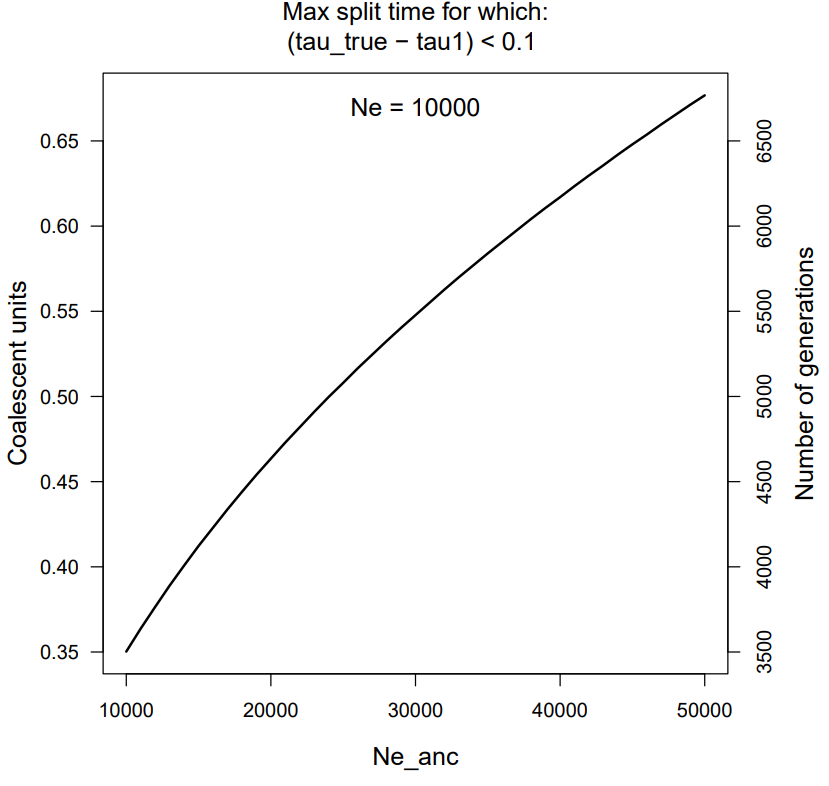

**Figure S2. When is the assumption of absence of novel mutations valid, such that can *τ* be reliably inferred from *F_ST_*?** Line graphs depicting maximum split times for which the true *τ*-value differs less than 10% from the estimate suggested by the formula: *τ* = ln(1-*F_ST_*)/(1000∙ln(0.999)). Note that the accuracy depends on the *N_e_* of extant populations, but also on the *N_e_* of the ancestral population. The reason is that small ancestral populations are expected to harbour low genetic diversity, meaning that for the same period of isolation a relatively higher proportion of *D_xy_* can be attributed to novel mutations, which violates the assumption that novel mutations are negligible.

SUPPLEMENTARY INFORMATION

**SI 1. Comparing Hudson *F_ST_* to Nei** (1973) ***F_ST_***

Nei (1973) estimated genetic distance between two populations as a function of probability of identity (i.e., homozygosity) within (*f_0_*) and between (*f_1_*) populations (SI2, SI3). Following Crow and Aoki (1984), these two parameters, which are the inverses of *π_xy_* and *D_xy_*, can be used to reformulate *F_ST_* (‘G_ST_’) as follows (Cockerham and Weir 1993; Slatkin 1993):

*F_ST_* = (*f_0_* – $\overline{f}$) / (1 – $\overline{f}$)

In which:

*f_0_*	= 1 – *π_xy_*	# probability of homozygosity within populations

*f_1_*	= 1 – *D_xy_*	# probability of homozygosity between populations

$\overline{f}$	= (*f_1_* + *f_0_*)/2	# probability of homozygosity in metapopulation

Rewriting, we can infer:

*F_ST_*	= (*f_0_* – $\overline{f}$)/(1-$\overline{f}$)

*F_ST_*	= (*f_0_* – (*f_1_* + *f_0_*)/2)/ (1 – (f_1_ + f_0_)/2)

*F_ST_*	= (*f_0_* – *f_1_*)/(2 – f_0_ – f_1_)

*F_ST_*	= ((1 - *π_xy_*) – (1 – *D_xy_*))/(2 – (1 - *π_xy_*) – (1 – *D_xy_*))

*F_ST_*	= (*D_xy_* – *π_xy_*)/ (*D_xy_* + *π_xy_*)

Thus, by rewriting this formula in terms of *π_xy_* and *D_xy_*, it can be shown that this metric is inconsistent with the *F_ST_*-metric of Hudson et al. (1992). Note that this outcome depends on the assumption $\overline{f}$= (f_1_ + f_0_)/2. Deriving $\overline{f}$from metapopulation allele frequencies (rather than by averaging of population averages) may produce a different outcome in case of unequal sample sizes.

**SI 2. Comparing Hudson *F_ST_* to Nei** (1977) ***F_ST_***

Nei (1977) defined *F_ST_* as:

*F_ST_*	=	(*H_T_* – *H_S_*) / *H_T_*

In which:

*H_T_*	= ½∙(*π_xy_* + *D_xy_*)	# probability of heterozygosity in metapopulation

*H_S_*	= *π_xy_*	# probability of heterozygosity within populations

Rewriting, we can infer:

*F_ST_*	=	(*H_T_* – *H_S_*) / *H_T_*

=	(½∙ (*π_xy_* + *D_xy_*) – *π_xy_*) / (½∙(*π_xy_* + *D_xy_*))

=	(-½*π_xy_* + ½*D_xy_*) / (½∙(*π_xy_* + *D_xy_*))

=	½(-*π_xy_* + *D_xy_*) / (½∙(*π_xy_* + *D_xy_*))

=	(-*π_xy_* + *D_xy_*) / (*π_xy_* + *D_xy_*)

=	(*D_xy_* – *π_xy_*) / (*π_xy_* + *D_xy_*)

Thus, by rewriting this formula in terms of *π_xy_* and *D_xy_*, it can be shown that this metric is inconsistent with Hudson *F_ST_*, but consistent with the *F_ST_*-metric of Nei (1977). Note that this outcome depends on the assumption *H_T_* = ½∙(*π_xy_* + *D_xy_*) (Nei 1973). Deriving *H_T_* from metapopulation allele frequencies (rather than by averaging of population averages) may produce a different outcome in case of unequal sample sizes.

**SI 3. Comparing Hudson *F_ST_* to Nei’s *D*** (Nei 1972)

Nei’s D, introduced by Nei (1972), defines genetic distance (*D*) and identity (*I*) for a pair of populations *X* and *Y* as:

D = -ln(*I*)

*I* = *J_xy_*/√(*J_x_***J_y_*)

In which:

*J_x_*	= 1 – *π_x_*	# probability of homozygosity within population *X*

*J_y_*	= 1 – *π_y_*	# probability of homozygosity within population *Y*

*J_xy_*	= 1 – *D_xy_* = *f_1_*	# probability of homozygosity between populations *X* and *Y*

When calculating Nei’s *D* from an unbiased selection of monomorphic and polymorphic sites, such that *J_XY_* is close to 1 (e.g. *J_XY_* for polar bears versus brown bears is 0.9967 (de Jong et al. 2023)), Nei’s *D* is an estimator of net divergence (*D_a_*) (Fig. S1):

Nei’s D	= *D_a_*

= *D_xy_* – *π_xy_*

If, instead, Nei’s *D* is calculated from a biased selection of monomorphic and polymorphic sites (e.g., SNPs only), this does not apply, and in that case Nei’s *D* returns an overestimate of *D_a_*.

**SI 4. Linear properties of the *f^2^*-statistic**

Net divergence, *D_a_*, the numerator of Hudson *F_ST_* (i.e., *D_xy_* – *π_xy_*), is equivalent to the *f^2^*-statistic, the numerator of Wright *F_ST_*. This *f^2^*-statistic, which rewrites *D_a_* as the squared difference in allele frequencies (*p*) in populations *X* and *Y*, *p_x_* and *p_y_* (Box 2), has useful linear properties which facilitates drift calculations.

In the absence of novel mutations, *p_x_* and *p_y_* are expected to remain on average equal to the allele frequencies in the ancestral population (*p_anc_*) and consequently also to each other:

*p_anc_*	= $\overline{p}$*_x_*	= $\overline{p}$*_y_*

Thus, the mean allele frequency difference (∆*p*) between two sister populations remains zero, no matter the magnitude of genetic drift (Fig. 2). If it was not for squaring ∆*p*, the *f^2^*-statistic would always reduce to zero.

Instead of altering the mean of ∆*p*, genetic drift affects the variation of ∆*p* across sites. Directly after a population split, little variation is expected in ∆*p* across sites. With time, random allele frequency fluctuations cause some sites to retain in both populations the same allele, while for other sites retain in either population different alleles. This increases variation in ∆*p* across sites, leading ultimately to a bimodal distribution in which for some sites ∆*p* equals 0 and for others ∆*p* equals 1 (Fig. 2). The *f^2^*-statistic, or between-population variance, quantifies this variation across sites, by estimating the mean per-site squared deviation in ∆*p* from the mean difference (which is zero).

The key to understanding that *F_ST_* is not affected by unequal population sizes, is the ‘second theorem of statistics’: the sum of variance. This theorem predicts that the variance of ∆*p* between any two populations *X* and *Y* is the sum of the within-population variance of allele frequencies and their covariances:

*f^2^	=*	Var(*p*)

*f^2^*	=	Var(*p_x_* – *p_y_*)

*f^2^	=*	Var(*p_x_*) + Var(*p_y_*) + Cov(*p_x_*,*p_y_*)

In the absence of gene flow, the fluctuations of allele frequencies in populations *X* and *Y* are completely unrelated, such that the covariance equals zero (i.e., Cov(*p_x_*,*p_y_*) = 0), and thus the formula simplifies to:

*f^2^	=*	Var(p_x_) + Var(p_y_)

=	1/L ∑ *(*p – $\overline{p}$*_x_*)^2^ + 1/L ∑ *(*p – $\overline{p}$*_x_*)^2^

As a comparison, when throwing one dice, the expected value is 3.5, with an expected variance of 2.92, implying that if this experiment is repeated many times, observed values will on average deviate from the mean by 1.71 (i.e., √2.92). (This indeed approximates the value obtained with the formula: ((4-3.5)+(5-3.5)+(6-3.5))/3.) When summing the outcome of two dice rolls, both the expected value and variance are doubled, equalling 7 and 5.84, respectively, with a range between 2 and 12. When scoring the difference between the two dice rolls – as in the case of allele frequency differences between a pair of populations – the expected value is 0 and the expected range is -5 to 5, but the variance is still 5.84.

The additivity of variance also applies to the accumulation of variance over generations. Thus, the expected variance of allele frequencies within populations at any point in time is, in principle, a linear function of effective population size (*Ne*) and population split time (*T*) (Fig. 2):

Var(p_x_)	=	(*p_x_**(1-*p_x_*))/(2*N_e_x_*) * T

And thus, for two populations combined:

*f^2^	=*	Var(*p_x_*) + Var(*p_y_*)

=	(*p_x_**(1-*p_x_*))/(2*N_e_x_*) * T + (*p_y_**(1-*p_y_*))/(2*N_e_y_*) * T

Because allele frequencies are bounded by the extreme values of 0 and 1, this linearity only holds when allele frequencies have not yet approached loss or fixation (Fig. 2). In any case, because of its linear properties, the sum of Var(*p_x_*) + Var(*p_y_*) will always equal the sum obtained if the variance within either population equaled the average variance of the two populations.

**SI 5. Different definitions/interpretations of net divergence**

We may distinguish between two different definitions of net divergence, namely:

*D_a_* = *D_xy_* – *π_xy_*

*D‘_a_* = *D_xy_* – *π_anc_*

*D’_a_* is the measure of interest if the objective is to quantify the proportion of novel mutations, which is proportional to split time. Unfortunately, *π_anc_* is typically unknown, and hence we use *D_a_* instead, with *π_xy_* replacing *π_anc_*.

*π_xy_* is usually defined as the mean nucleotide diversity averaged over the two populations (ref):

*π_xy_ = ½(π_x_ + π_y_)*	# common definition of *π_xy_*

*π_xy_* is an unreliable proxy of *π_anc_*, which can be overestimate in case of recent population size expansion, and an underestimate (more typically) in case of a recent population size decrease. Especially on short time scales, when novel mutations are negligible while genetic drift may not, *π_anc_* sets the upper limit of *π_xy_*. If the objective is to discern split times, a more accurate proxy of *π_anc_* is obtained by selecting the higher *π*-estimate among a pair of populations:

*π_xy_ = max(c(π_x_,π_y_))*

**SI 6. Two approaches for translating *F_ST_* into coalescent units (*τ*)**

Consider two populations, *A* and *B*, with an absolute genetic distance of 0.004, and a mean nucleotide diversity of 0.001, implying: *F_ST_* = (0.004-0.001)/0.004 = 0.75. Assuming that the difference between *D_xy_* and *π_xy_* is due solely to a reduction of *π_xy_* owing to genetic drift, we can infer the split time in terms of coalescent units (*τ*, i.e., multiples of 2*N_e_*), as follows:

*τ_π_*	= ln(1-*F_ST_*) / (1000∙ln(0.999))	= ln(1-0.75) / (1000∙ln(0.999))	= 1.3862

If assuming, instead, population size constancy, such that the difference between *D_xy_* and *π_XY_* is due solely to an increase of *D_xy_* owing to novel mutations, we infer:

*τ_D_*	= *F_ST_* /(1-*F_ST_*)	= 0.75/(1 – 0.75)	= 3

These estimates are normally incongruent, but in one specific condition they produce the same outcome. For the second scenario, *π_xy_* is constant and equals 0.001. Assuming equilibrium such that *π* = 4*N_e_µ*, and assuming a per-generation mutation rate (*µ*) of 10^-8^, Ne equals 25.000. We obtain a split time estimate of (*τ_D_*∙2∙25000 =) 150000 generations. Indeed, we find: *D_xy_* = 0.001 + 150000 ∙ 2 ∙ 10^-8^ = 0.004.

In the former scenario, *π_xy_* decreases exponentially from 0.004 to 0.001, with a mean value of 0.002161, implying a mean *N_e_* of approximately 54025 if assuming *π* = 4*N_e_µ*. This returns a split time estimate of (*τ_π_*∙2∙54025 =) 149779 generations. Indeed, we find: *π_xy_* = 0.004 ∙ (1- 1/(2∙54025))^149779 = 0.001. (Of course, this is an extremely unrealistic scenario, as 149779 generations would allow for plenty of novel mutations.)

**SI 7. R scripts for simulations**

\# Function to predict *D_xy_*, *π_xy_*, *F_ST_*, and *τ* as an iterative function of *N_e_*, *T* and *μ*:

predictfst<-function(ngen=50000,Ne=10000,u=10^-8,Ne_anc=10000)

{

\# u	= mutation rate per generation

\# Ne	= mean effective population size of two extant sister populations

\# Ne_anc	= effective population size of ancestral population, prior to split event

k	<- 1-(1/(2*Ne))	# drift factor (i.e., amount of variation retained)

pi_anc	<- 4*Ne_anc*u	# ancestral nucleotide diversity (assuming drift-mutation equilibrium)

dxy	<- pi_anc	# at T = 0, Dxy equals pi_anc

pixy	<- pi_anc	# at T = 0, pi equals pi_anc

for(n in c(2:ngen))

{

\# infer Hudson Fst:

pixy	<- k*(pixy)+2*u	# pixy at generation n

dxy	<- dxy+2*u	# dxy at generation n

fst	<- (dxy-pixy)/dxy	# fst at generation n

\# infer coalescent units (number of generations divided by 2Ne):

tau_true	<- n/(2*Ne)	# true estimate

tau_1	<- log(1-fst)/(1000*log(0.999))	# assuming delta_Dxy=0

tau_2	<- fst/(1-fst)	# assuming delta_pixy=0

}

}

\# Function to predict *D_xy_*, *π_xy_*, *F_ST_* and *Nei’s D* by simulating allele frequency fluctuations (assuming the absence of novel mutations):

driftsim<-function(nloci=100000,ne1=1000,ne2=1000,p=0.5,ngen=750)

{

\# generation 1 (split event, both populations start with same ancestral variation):

pop1	<- rpois(n=nloci,lambda=p*100)/100

pop2	<- pop1

for(n in c(2:ngen))

{

\# single-locus estimates (vectors):

pop1	<- rbinom(prob=pop1,size=2*ne1,n=nloci)/(2*ne1)

pop2	<- rbinom(prob=pop2,size=2*ne2,n=nloci)/(2*ne2)

metapop	<- (pop1+pop2)/2	# allele frequency in metapopulation

\#

\# multi-locus summary statistics (scalars):

p1	<- mean(pop1)

p2	<- mean(pop2)

pmeta	<- mean(metapop)

dxy	<- mean(pop1*(1-pop2)+(1-pop1)*(pop2))

pi1	<- mean(2*pop1*(1-pop1))

pi2	<- mean(2*pop2*(1-pop2))

pixy	<- (pi1 + pi2)/2

f2	<- mean((pop1-pop2)^2)

f1	<- mean(pop1*pop2+(1-pop1)*(1-pop2))

f0_0	<- mean(pop1^2+(1-pop1)^2)

f0_1	<- mean(pop2^2+(1-pop2)^2)

f0	<- (f0_1+f0_2)/2

\#

\# Fst, Dxy and Da:

hudsonfst	<- (dxy-pixy)/dxy	# Hudson et al (1992) Fst

crowaokifst	<- (f0-f1)/(1-f1)	# Crow and Aoki (1984) Fst

wrightfst	<- f2/(2*pmeta*(1-pmeta))	# Wright (1943) Fst

neiI	<- f1/sqrt(f0_1*f0_2)	# Nei’s I

neiD	<- -log(neiI)	# Nei’s D

}

}
